# A note on dynamical compensation and its relation to parameter identifiability

**DOI:** 10.1101/123489

**Authors:** Omer Karin, Uri Alon, Eduardo Sontag

## Abstract

We recently identified a motif for dynamical compensation (DC) – a property where a system maintains the dynamics and steady-state of a regulated variable robust in the face of fluctuations in key parameters. Such parameters are therefore unidentifiable from measurements of the regulated variable at steady-state. On the other hand, since the models showing dynamical compensation are typically non-redundant, their parameters are identifiable from experimental data. We clarify this apparent discrepancy by requiring that the parameters of DC circuits be identifiable both away from steady-state and when measuring other system variables. We use this observation to provide a definition for DC in terms of parameter identifiability and discuss its relevance for the examples provided in Karin et al.

## Introduction

Biological systems often need to maintain their steady-state output robust in the face of variation, a property known as exact adaptation [1]. Motifs for exact adaptation include integral feedback loops [2]. Some biological systems must go beyond maintaining the steady state output, and to keep their entire dynamical trace, including amplitude and response time, robust to fluctuations in key parameters. We called this robustness of the entire trace ‘dynamical compensation’ (DC) [3].

We recently identified a motif for DC in physiological circuits, including endocrine circuits [3]. In this motif, a regulated variable directly controls the functional mass of its regulating tissue. This motif creates a feedback loop on slow timescales that adjusts the functional mass of a tissue that, for example, secretes a hormone, to compensate for changes in parameters such as the hormone sensitivity of the target tissues. This motif occurs in the glucose-insulin system, and may apply to several other physiological circuits, including the regulation of calcium by PTH, the regulation of arterial oxygen by the carotid body, and neuroendocrine circuits [3].

The original paper described DC as follows. Consider a system with an input *u(t)* and an output *y(t*,*p)* such that *p* > *0* is a parameter of the system. The system is initially at steady state with *u(0)* = *0*. DC with respect to *p* is the property that for any input *u(t)* and any (constant) *p* the output of the system *y(t*,*p)* does not depend on *p*. That is, for any *p*,*q* and for any time-dependent input *u(t), y(t*,*p)* = *y(t*,*q)*. Although the paper also dealt with non-steady-state situations, this description suggests that the parameter *p* is unidentifiable from measurements of *y*.

Identifiability is a concept from structural identifiability (SI) theory, a mathematical theory that deals with unidentifiable parameters – parameters that cannot be fit from experimental data [4-5]. Such parameters are often considered redundant. SI theory is therefore useful for finding whether a model is well defined, or whether it should be reformulated with less parameters. Since DC models have parameters that cannot be identified from measurements of their regulated parameter, it has been suggested that SI theory may be useful for finding models with the DC property, as pointed out in [6-7].

Although it was not explicitly stated in Karin et al., the examples given in that paper all have an additional feature, namely that the DC property (a robustness property) must, in addition, be non-redundant. This creates an apparent discrepancy, which can be resolved by tightening the definition to explicitly require that while the parameter *p* of a DC model is unidentifiable from measurements of *y* at steady-state, it should be identifiable from other experimental measurements – either from measurements of *y* away from steady-state or from measurements of other system variables. We use this intuition to refine the definition of DC using SI theory.

Informally, we will say that a model has DC on a variable *y* with respect to a parameter *p* if the model is well defined (i.e *p* is identifiable from experimental data if rich enough perturbations of parameters are allowed and/or rich enough types of measurements are allowed) but the system has Structural Non Identifiability (SNI) for *p* when only measuring the variable *y* starting from steady-state. We next define this concept formally.

### Definition

Consider a dynamical system that is defined as follows:

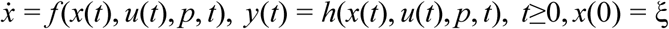

Where *f, h* describe respectively the dynamics and the read-out map; *u* = *u(t)* is an input (stimulus, excitation) function, assumed to be piecewise continuous in time, *x(t)* is an n-dimensional vector of state variables, *y(t)* is the output (response, reporter) variable, and *p* is the parameter (or vector of parameters) that we wish to focus our attention on. ξ is the initial state of the system. We require that when the input *u*=*0* this system should have a unique steady-state *x*_*p*_, i.e a unique solution for *f (x(t)*, 0,*p)* = 0.

For the system to show DC with respect to *p* it must show both [I,II] and either [III] or [IV]:

I. Exact adaptation with respect to changes in the parameter *p*, that is, for all parameters *p* and all initial conditions ξ then *h(x*, 0,*p)* → *y*_0_.
II. SNI, meaning that for all *p*,*q*, for all inputs *u(.)*, for all t, *y(x*_*p*_,*p, u, t)* = *y(x*_*q*_, *q, u, t)*
III. Identifiability from perturbations: For all *p≠q*, there is some input *u(.)* and some time *t*_*0*_ s.t. *y(xp, q, u, t*_0_) *≠ y(xq, q, u, t*_0_).

If there exists an additional output function:

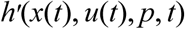

then the model may have DC if [I,II] hold when only the output *h* is available, and also:

Complete identifiability given both *h*,*h’:* for any *p≠q* there exists *u(.)* and a *t*_*0*_ s.t.

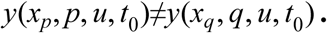

## Parameter identifiability of a DC model

Here we apply the above definition to the *βIG* model for the glucose-insulin-beta cell system from Karin et al [3]. The system is defined by the following equations:

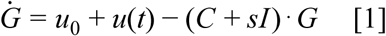

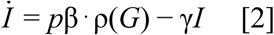

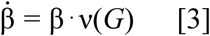
where *G* is plasma glucose concentration, *I* is plasma insulin concentration, *u*_0_ is endogenous production of glucose, *u(t)* is meal intake, *C* is glucose removal rate at zero insulin and *s* is insulin sensitivity. Secretion of insulin is proportional to beta cell functional mass *β*, where ρ(*G*) is a monotonically increasing function of *G*, γ is the insulin removal rate, and *p* is the insulin secretion per cell. Finally, ν is the beta cell growth rate and it is stable at the homeostatic glucose set point *G*=*G*_*0*_, that is v(*G*_0_) = 0.

As was shown in Karin et al. [3], this system has both exact adaptation and the parameters *s*,*p* are unidentifiable from measurements of *G* at steady-state, and therefore conditions [I-II] hold. We now show that *s*,*p* are identifiable from perturbations or, alternatively, from measurements of additional variables.

We first calculate the steady-state of the system for a given set of parameters *s, p* and assuming zero input *u* = 0. Because the system has exact adaptation, steady-state glucose is constant *G*_*ST*_ = *G*_0_. The steady-state level of insulin is then:

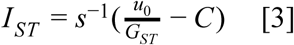
and the steady-state beta cell functional mass is:

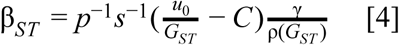
By measuring [3,4] we can infer both *s, p*, so condition [IV] holds given that we can measure both insulin and beta cell functional mass.

The parameters can also be identified by measurements of glucose off steady-state. Let the system have insulin secretion *p* and insulin sensitivity *s*_1_ that changes from *s*_1_→*s*_2_ (Figure 1). Just after the change *s*_1_→*s*_2_ the level of glucose changes to some level *G*≠*G*_*ST*_ and returns to G = *G*_*ST*_ only after a transient period. During the transient period, the dynamics of glucose in response to inputs is different from its dynamics before the change *s*_1_→*s*_2_ and specifically its response to zero input *u*=*0* is different.

**Figure 1.**
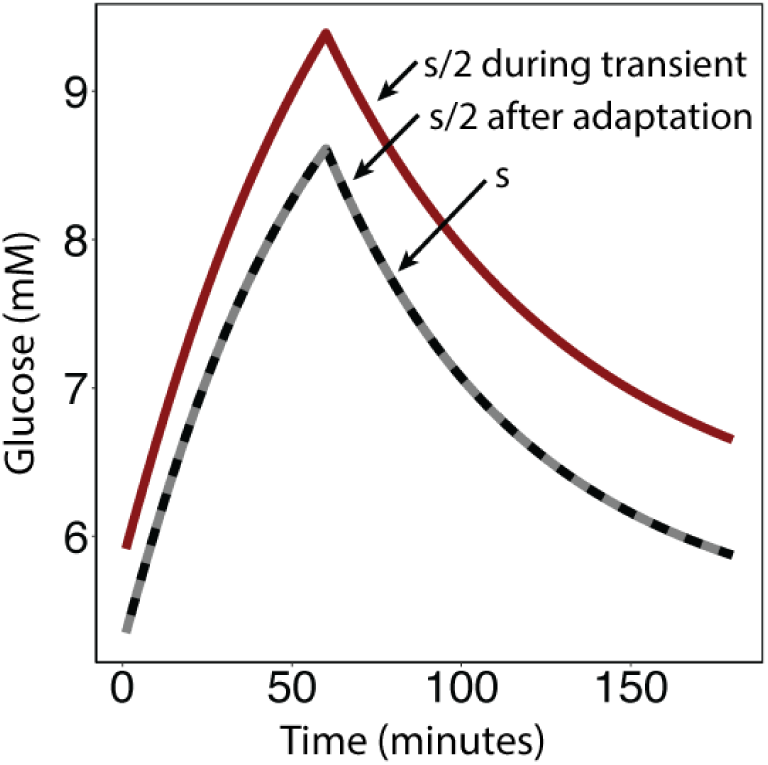
Dynamical compensation is only guaranteed after adaptation. Glucose dynamics in response to meal before a change in insulin sensitivity s→s/2 (black), immediately after the change in insulin sensitivity (red) and after adaptation (dashed gray). Glucose dynamics are the same before the change in insulin sensitivity and after adaptation. However, during the transient after the change in insulin sensitivity, and before adaptation, glucose steady-state and dynamics are perturbed.

The value of *s*_2_ can be approximated after the step change *s*_1_→*s*_2_, assuming that insulin changes much faster than beta cell functional mass. Let 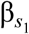 be the steady-state beta cell functional mass when insulin sensitivity *s* = *s*_1_. After the change *s*_1_→*s*_2_ glucose changes to some value *G*′≠*G*_*ST*_ and insulin changes to some value *I*′ so 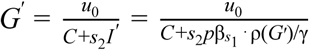. Since ρ(*G*′) is monotonic and *G* and the other variables are non-negative then this equation has a unique solution for each *s*_2_ and therefore the input *u*=*0* suffices to distinguish between each 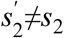. Therefore given only measurements of β one can infer s after any perturbation (Figure 1).

Condition [III] thus holds with respect to s, given knowledge of *p*. When *p* is not known then we require an additional measurement (e.g of beta cell mass at steady state/off steady state). This is a plausible measurement of a physiological quantity. Thus, given *p* we can infer s either from either (i) measurements of glucose and insulin at steady state, or (ii) measurements of glucose off steady state. To infer *p* we only require some additional measurement such as beta cell mass.

To conclude, DC is a robustness property where the dynamics of a regulated variable are robust to fluctuations in key parameters. This robustness is only guaranteed at steady-state. This is the origin of the name dynamic compensation: when a parameter changes, the system slowly adjusts to precisely compensate for the change, and full compensation is achieved at steady state. DC models must be non-redundant, so their parameters can be fit either from measurements away from steady-state or from measurements of other system variables.

